# Mutation accumulation differentially impacts ageing in mammalian tissues

**DOI:** 10.1101/247700

**Authors:** Zeliha Gözde Turan, Poorya Parvizi, Handan Melike Dönertaş, Jenny Tung, Philipp Khaitovich, Mehmet Somel

## Abstract

Medawar’s mutation accumulation (MA) hypothesis explains ageing by the declining force of natural selection with age: slightly deleterious germline mutations that are functional in old age are not effectively eliminated by selection and therefore lead to ageing-related phenotypes. Although widely cited, empirical support for the MA hypothesis, particularly molecular evidence, has remained limited. Here we test one of its predictions, that genes relatively highly expressed in old adults vs. young adults should be under weaker purifying selection than those relatively highly expressed in young adults. To do so, we combine 23 RNA-sequencing and 35 microarray gene expression datasets (including 9 tissues from 5 mammalian species) with protein and regulatory sequence conservation estimates across mammals. We identify age-related decrease in transcriptome conservation (ADICT) in four tissues, brain, liver, lung, and artery, but not in other tissues, most notably muscle and heart. ADICT is driven both by decreased expression of highly conserved genes and up-regulation of poorly conserved genes during ageing, in line with the MA hypothesis. Lowly conserved and up-regulated genes in ADICT-associated tissues have overlapping functional properties, particularly involving apoptosis and inflammation, with no evidence for a history of positive selection. Our results suggest that tissues vary in how evolution has shaped their ageing patterns. We find that in some tissues, genes up-regulated during ageing, possibly in response to accumulating cellular and histological damage, are under weaker purifying selection than other genes. We propose that accumulation of slightly deleterious substitutions in these genes may underlie their suboptimal regulation and activity during ageing, shaping senescent phenotypes such as inflammaging.

## Introduction

To date, more than 300 theories have been postulated to explain senescence, *i.e.* age-related loss of function and increase in mortality rates^1^. The mutation accumulation (MA) hypothesis, an evolutionary explanation for ageing first developed by J.B.S. Haldane^2^ and Peter Medawar^3^, is among the most influential of such theories. It states that negative selection against germ-line substitutions that exhibit harmful effects only during old age will be inefficient. Such substitutions can eventually fix through genetic drift, and thus contribute to observed senescent phenotypes^4^. The MA hypothesis generates several testable predictions. One expectation is that genetic variance in fitness-related traits, such as reproductive success or survival, will increase with age^2,5^. A second expectation is that inbreeding depression will also increase with age. In line with these predictions, several studies have shown age-related increase in genetic variance in fitness-related traits in either laboratory (*Drosophila melanogaster:* refs. ^6,7^) [but see refs. ^8–10^] or natural populations (Soay sheep, red deer: ref. ^11^). In humans, heritability of CpG methylation patterns was shown to increase with age for about 100 genome-wide loci also consistent with MA, although possible fitness consequences were not evaluated^12^. In an indirect test of the expectation for inbreeding depression, outbreeding was reported to reduce age-related increase in mortality in hermaphroditic snails^13^, again in line with MA. Finally, Rodríguez et al.^14^ studied >2,500 human genetic variants linked to 120 genetic diseases and reported that variants associated with late-onset disease segregate at higher frequencies than those associated with early-onset disease, a third prediction under MA^14^.

Beyond those cited above, relatively few studies have used empirical data to test the MA hypothesis. In particular, the possibly variable contribution of MA in the ageing processes affecting different species and their different tissues has not yet been tackled. In addition, we have limited understanding of the nature and prevalence of late-expressed substitutions, with the exception of a few extreme cases such as the CAG repeat variants in the *huntingtin* gene that cause Huntington's disease^5^.

The role of MA in ageing therefore awaits testing through new approaches that encompass a larger number of traits, a wider array of species, different tissues, and that include molecular data. One such approach would be to take advantage of widely available transcriptome data, in particular genome-wide gene expression datasets that comprise adult individuals of varying age. Such transcriptome datasets have traditionally been used to identify functional processes affected by or underlying senescence, although they can also be used to test evolutionary theories, as we show here.

In previous work, we used prefrontal cortex transcriptome age-series from humans to test whether protein sequence conservation varies among genes that are highly expressed at different ages^15^. This analysis showed that are relatively highly expressed genes in young adults vs. old adults are evolutionarily more conserved than those that are relatively highly expressed genes in old adults vs. young adults, which we call <u>a</u>ge-related <u>d</u>ecrease <u>i</u>n <u>c</u>onservation of the <u>t</u>ranscriptome (ADICT), consistent with the MA hypothesis. However, this work analysed only one brain region and did not distinguish between two distinct processes: (a) up-regulation of lowly conserved genes with age and, (b) down-regulation of highly conserved genes with age. Both processes could cause the ADICT effect, but only (a) would be predicted by MA.

To address these limitations, here we expanded our analysis to include 9 different tissue types and 5 mammalian species. First, we investigated the prevalence of the ADICT pattern across multiple mammalian ageing datasets, using estimates of protein and regulatory sequence conservation across mammals. Second, we determined whether genes up-regulated late in life show low evolutionary conservation, as predicted by MA. In other words, we tested whether slightly deleterious mutations are more likely to fix in genes that are more highly expressed in old age, such as genes that respond to age-associated tissue damage^16,17^.

## Results

### Age-related decrease in conservation of the transcriptome

We collected published transcriptome age-series of young and old adults of 5 mammalian species, generated using RNA-sequencing or microarrays (*Homo sapiens, Macaca mulatta, Macata fascicularis, Rattus norvegicus, Mus musculus; n* = 58 datasets and 2041 unique samples in all). The datasets include different brain regions (humans, macaques, rats, and mice), muscle (humans, rats, and mice), artery (humans, macaques, and rats), heart (humans), skin (humans and mice), kidney (humans and mice), liver (humans and mice), lung (humans and mice), and spleen (mice). Across all analysed datasets (*n* = 9 to 116 individuals, mean = 35.2), human ages range from 16 to 106 years, macaque ages range from 4 to 28 years, rat ages range from 3 to 30 months, and mouse ages range from 8 to 130 weeks (Supplementary Table 1).

We studied congruence in age-related gene expression change across the 58 datasets. First, for each gene in each dataset we calculated the Spearman correlation coefficient between gene expression level and individual age (*ρ_EA_*). We then compared datasets to estimate pairwise similarity in *ρ_EA_* values across common genes. *ρ_EA_* values were mostly (72% of comparisons) positively correlated across data sets, indicating that the same genes’ expression levels were similarly affected by age (Supplementary Fig. 1).

As measure of gene sequence conservation we used estimates of purifying selection on protein sequence (ω_0_), calculated by Kryuchkova-Mostacci and Robinson-Rechavi (2015)^18^ and estimated for the human or the mouse branch using the branch-site model^19^. ω_0_ is the *dN/dS* ratio calculated for those sites determined to be under purifying selection, and thus is expected to be a direct measure of the strength of purifying selection on a gene. We further calculated an adjusted protein conservation metric (ω_0_^*^) for each gene, factoring out the possible effects of GC content, CDS length, intron length, intron number, mean expression, median expression, maximum expression, tissue specificity, network connectivity, phyletic age, and number of paralogs, using a multiple regression model following Kryuchkova-Mostacci and Robinson-Rechavi (2015)^18^. The value -ω_0_^*^ (ω_0_^*^ multiplied by -1) represents the main protein sequence conservation metric we used in our analyses, where more positive values represent more conserved genes.

We then investigated the prevalence of ADICT in mammalian ageing. To do so, we first calculated the Spearman correlation coefficient between gene expression level and the protein sequence conservation metric (which we call *ρ_EC_*) across all genes, for each individual in each dataset (Figs. 1a, 1b). Note that the conservation metric (-ω_0_^*^) is a constant value per gene, while gene expression levels will differ among individuals. In general, a positive *ρ_EC_* is expected, such that more highly expressed genes tend to be more conserved in their protein sequence^20^, but its degree may vary among individuals depending on age. To test this idea, in each dataset we determined the correlation between individual ages and *ρ_EC_* (*ρ_A_ρ_EC_*). Fig. 1c provides an example of such a pattern in one brain ageing dataset^21^, and Fig. 2 shows the results across all datasets. The relation between individual ages and *ρ_EC_* was found to follow a mainly linear trajectory (Supplementary Fig. 9 and Supplementary Table 3).

**Fig. 1.**
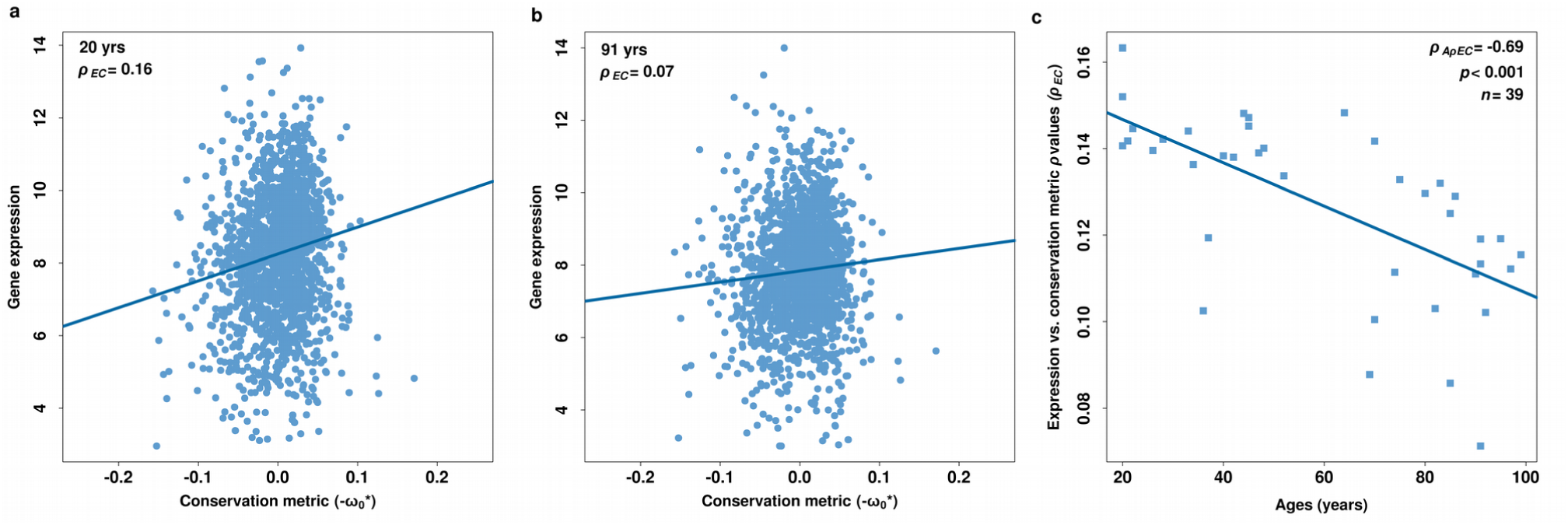
Relationship between gene expression level and protein conservation. Examples of gene expression level - protein conservation metric correlations **(a)** for a 20 year-old human, and **(b)** for a 91-year old human, in the postcentral gyrus of the brain (data from ref. ^21^). The analysis includes only age-related genes detected in this dataset (at *q* < 0.10). Each point represents a gene (*n* = 1688). The x-axis shows the protein sequence conservation metric, where more positive values reflect higher conservation across mammals. The y-axis shows log2 transformed gene expression level. The expression-conservation *ρ* values (*ρ_EC_*) are indicated in the inset. To improve visualization, we removed genes with very low conservation metrics (*n* = 3) in panels (a) and (b). Note that our correlation statistic, Spearman, is not affected by such potential outliers. **(c)** Age-dependent change in expression-conservation *ρ* values in the human postcentral gyrus, based on age-related genes in the same dataset as panels (a) and (b). The y-axis shows expression-conservation *ρ* values (*ρ_EC_*) calculated for each individual in this dataset (*n* = 39). The x-axis shows the age of an individual. The *ρ* value between age and expression-conservation correlation (*ρ_A_ρ_Ec_*) is indicated in the inset.

**Fig. 2.**
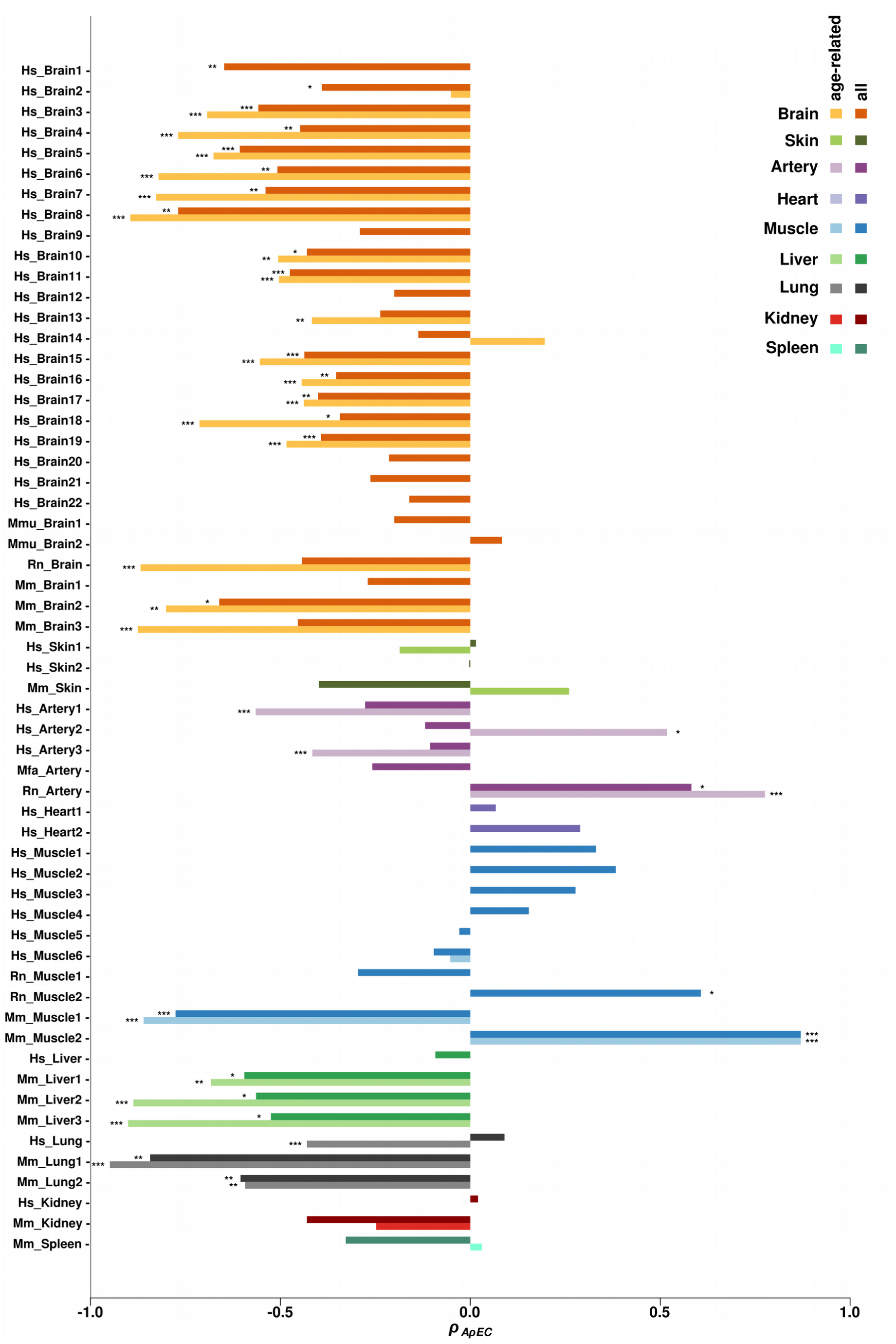
Age-dependent changes in transcriptome conservation. The x-axis shows the Spearman correlation coefficient (*ρ_A_ρ_Ec_*) between individual age and expression-conservation correlations (*ρ_Ec_* described in Fig. 1). The statistics are calculated separately for each dataset, and for significant age-related genes in that dataset (light bars), as well as for all expressed genes (dark bars). Note that in 21 of 58 datasets (cases where the light bar is missing), no significant age-related gene set could be identified (at *q* < 0.10). The asterisks indicate nominal significance of the Spearman correlation test, (*): *p* ≤ 0.05, (**): *p* ≤ 0.01, (***): *p* ≤ 0.001. In the analysis using age-related genes, the 25 datasets showing nominal significance for ADICT remained significant at *q* < 0.10 after applying Benjamini-Hochberg correction (excluding one muscle dataset showing ADICT). In the analysis using all genes, the 21 / 22 datasets showing nominal significance for ADICT remained significant at *q* < 0.10 after applying Benjamini-Hochberg correction, with “Mm_Liver3” the only exception.

In each dataset, we used two gene sets for testing ADICT: (a) genes that showed significant age-related change in expression levels (at Spearman correlation test *q*-value < 0.10), and (b) all expressed genes. We conducted analyses using all expressed genes in order to avoid a reduction in statistical power in datasets with low sample sizes and to determine whether patterns that hold for strongly age-associated genes also apply across the entire transcriptome (Supplementary Table 2). Note that in 21 of 58 datasets, and in particular smaller datasets, we could not identify a set of significant age-related genes at *q* < 0.10; for these datasets we only conducted the analysis using all expressed genes (see Methods).

In the brain, for 18 / 19 datasets with age-related genes, we found negative *ρ_A_ρ_EC_* values consistent with the notion of ADICT. *ρ_A_ρ_EC_* values for 17 of these 18 datasets were significant at nominal *p* < 0.05. When repeating this analysis with all expressed genes, 27 / 28 datasets had negative *ρ_A_ρ_EC_* values, 16 of which were nominally significant. Together, these results support a general trend of ADICT in the brain (Fig. 2). We also found significant negative *ρ_A_ρ_EC_* values in the majority of liver (3 / 4) and lung (3 / 3) datasets, and in 2 of 5 artery datasets. In contrast, we found no consistent ADICT pattern in various muscle types (*n* = 10, where we identified only one nominally significant case) or in heart, skin, spleen or kidney. We also note that in 6 / 10 muscle datasets, we did not detect significant age-related expression change in gene-by-gene analyses (Supplementary Table 2). Overall, 26 / 58 of the datasets showed significant ADICT signatures across age-related genes after correction for multiple testing (*q* < 0.10), with this pattern driven by brain, liver, lung, and artery (Fig. 2). ADICT signatures were also consistent with our findings using -ω_0_^*^ when we used the mean PhastCons score as the measure of negative selection on protein coding sequences (Supplementary Fig. 7).

To determine the robustness of this result with respect to our protein-coding sequence conservation metric, we repeated the analysis using ω_0_ values without applying multiple regression, as well as ω values obtained from the Ensembl database for “one-to-one orthologs” between human-mouse, human-elephant, and human-cow (*i.e.* raw *dN/dS* values). We further tested whether the trend holds when we exclude (a) putatively positively selected genes (with ω >1 in our data), (b) immune system genes known to be generally fast evolving^19,22,23^, and (c) genes down-regulated with age in each dataset (ranging from *n* = 1,086 to 6,717). In addition, to exclude the possibility that ADICT signals are driven by gene expression changes involving only few functional processes (*e.g.* highly conserved developmental genes being down-regulated), we calculated *ρ_A_ρ_EC_* separately for genes in each of the largest GO Biological Process categories (*n* = 19, each with node size >1000 annotated genes) (Supplementary Fig. 2). Negative *ρ_A_ρ_EC_* values were repeatedly detected in the same 25 datasets (excluding one muscle dataset showing ADICT), irrespective of the metric used, the gene sets, and GO categories involved (Supplementary Table 2 and Supplementary Fig. 2). ADICT thus appears to be a consistent trend in multiple tissues, although it is also not a universal pattern across all mammalian tissue types.

### Distinct processes contribute to ADICT

We next investigated two non-exclusive scenarios that could lead to ADICT: (a) genes that show age-related increase in expression could be lowly conserved, consistent with MA, or (b) genes that show age-related decrease in expression could be highly conserved, relative to genes that show no change in expression. The latter scenario could occur if a set of highly conserved genes (*e.g.* synaptic genes) are down-regulated during the postnatal lifespan, as previously reported^15,24^, but would not provide direct evidence in support of MA.

To test which of these scenarios underlie ADICT, we compared the mean conservation metric among (a) genes that show increases in expression with age (*i.e.* up-regulation, *ρ_EA_* > 0.1, *q* < 0.1), and (b) genes that show decreases in expression with age (*ρ_EA_* < -0.1, *q* < 0.1), using (c) genes that show no age-related changes in expression level as a control. We repeated this analysis across the 25 datasets showing the ADICT signature. We found results consistent with both scenarios (Fig. 3): genes that show decreases in expression with age were more strongly conserved than genes with no change (*n* = 22 / 25; 14 with bootstrap support >95%). Genes up-regulated with age were also more weakly conserved than genes with no change, in nearly all cases (*n* = 23 / 25; 15 with bootstrap support >95%). The latter observation is in line with the MA hypothesis.

**Fig. 3.**
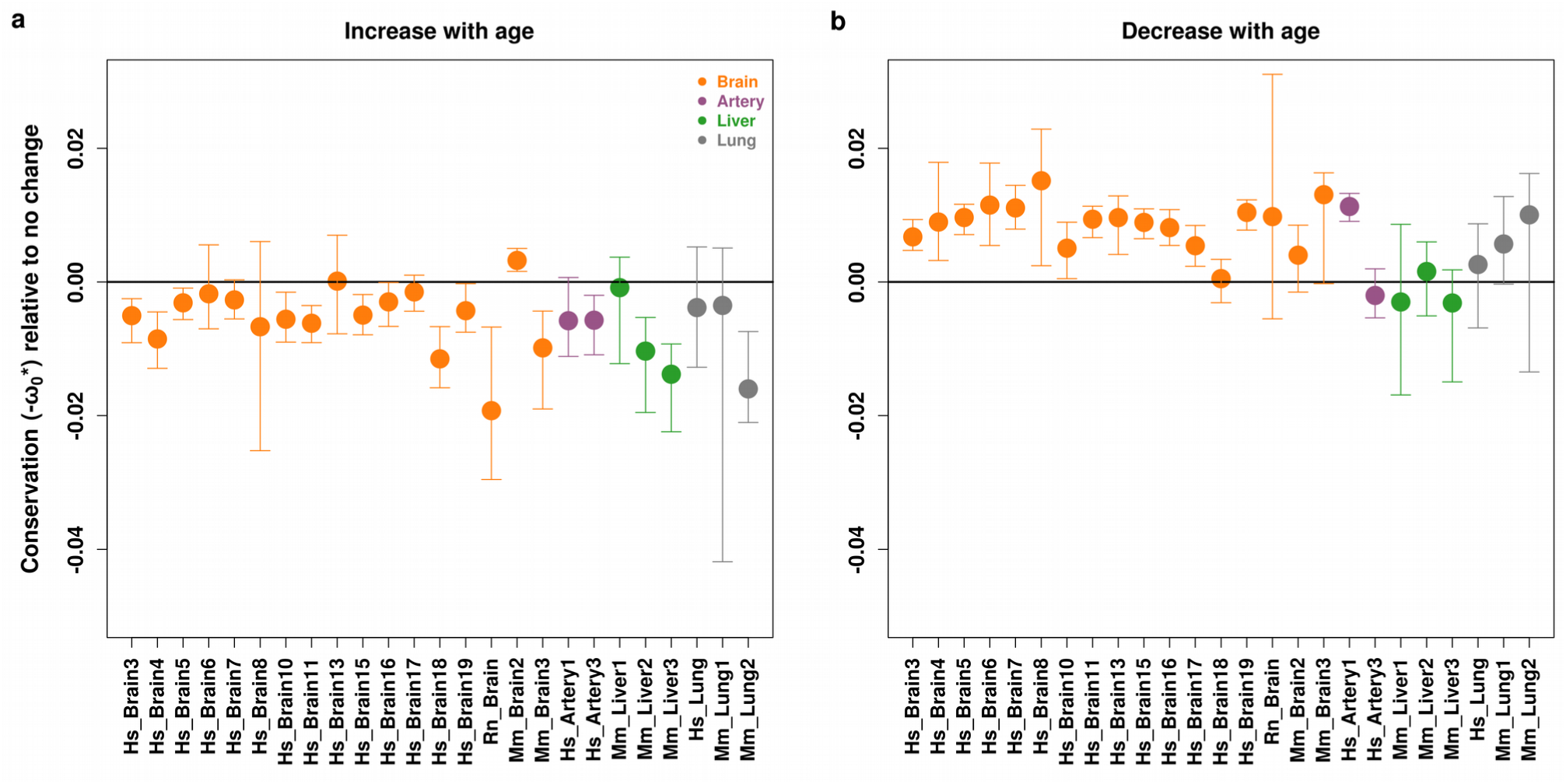
Mean conservation metrics among gene sets with different patterns of age-related change in expression levels. The plots show mean conservation metric for genes that show age-related increase (a) and age-related decrease (b) in expression levels, compared to mean conservation metric among genes that show no significant age-related change in expression level (see Methods). The error bars indicate 95% confidence intervals calculated by 1,000 bootstraps, for the 25 datasets consistent with ADICT.

We further hypothesized that genes that more frequently exhibit increased expression with age across tissues would be less likely to be conserved. To test this hypothesis, we selected genes shared across the 25 ADICT datasets and counted how many times each gene was up-regulated with age. As predicted, we observed a negative correlation between the number of cases in which a gene was up-regulated with age and its conservation metric (*ρ* = -0.17, *p* < 0.001) (Supplementary Fig. 8a). We further repeated this analysis for all 58 datasets, which also revealed a significant negative correlation (*ρ* = -0.24, *p* < 0.001) (Supplementary Fig. 8b).

### Functional analysis of ADICT

To find functionally coherent gene sets that may contribute to ADICT patterns in brain, liver, lung, and artery, we conducted Gene Ontology (GO) analysis for the 3 GO domains (Biological Process, Cellular Component, Molecular Function). We calculated statistics for GO analyses by ranking genes according to both the conservation metric (-ω_0_^*^) and expression-age correlations (*ρ_EA_*) and investigating GO term enrichment in the 10% tails of the distributions. We separately analysed (a) genes that showed increased expression with age and low relative conservation (IELC, consistent with MA), and (b) genes that showed decreased expression with age and high conservation (DEHC). We sought shared GO categories enriched in either IELC genes or DEHC genes across all 25 datasets showing the ADICT signature. To determine the random expectation for shared GO categories, we randomly permuted ages of individuals in each dataset 1000 times, calculated *ρ_EA_*> again, and repeated the gene ranking and GO analysis.

IELC genes were enriched in the same 24 GO Biological Process categories in all the 25 datasets (expected = 0; permutation test *p* < 0.001) (Supplementary Figs. 3 and 4, Supplementary Table 4). These included categories related to apoptosis, inflammation, and the immune response, among others (see the REVIGO summary in Supplementary Fig. 4). In addition, four GO Cellular Component categories (expected = 0; *p* < 0.001) and one GO Molecular Function category (expected = 0; *p* = 0.022) were shared among IELC genes across the 25 datasets (Supplementary Fig. 3). Among DEHC genes, we found shared enrichment only in Cellular Component and Molecular Function categories (permutation test *p* < 0.05); significant gene sets included synapse and signaling related functions (Supplementary Fig. 4 and Supplementary Table 4). Although functional analysis suggests DEHC as a potential factor for functional decline, we will only focus on IELC as the objective of this study is to test MA hypothesis.

### Age-dependent effects on regulatory region conservation

Finally, we asked whether ADICT also extended to conservation of transcriptional regulatory regions. To test this possibility, we calculated the mean PhastCons score (a metric for conserved elements) from the UCSC database based on a 100-way vertebrate alignment^25^, for (a) +/- 2000 bp around the transcription start site (TSS) and (b) the 3’-UTR. We then repeated our ADICT analysis by substituting these two regulatory conservation metrics for -ω_0_^*^. This again revealed a heterogeneous trend toward ADICT across tissues, with consistent ADICT trends in brain, liver, and lung (Supplementary Fig. 5 and Supplementary Fig. 6).

## Discussion

The MA hypothesis predicts that burden of slightly deleterious germline substitutions will increase with age due to the declining force of negative selection^3^. Our approach differs from earlier attempts to test this hypothesis^6–11,13,14^ in two respects. First, instead of relying on intra-species variation to estimate mutational load, we used inter-species divergence, which may be statistically more powerful as it involves a larger number of substitutions. Second, we studied the mutational load on multiple tissues, thus taking into account the possibility that age-dependent germline mutational load may vary among tissues, depending on tissue-specific developmental patterns, mitotic capacity, and consequences for organism-level fitness.

We found the age-related decrease in transcriptome conservation (ADICT) pattern in datasets from brain, liver, lung, and artery, consistently across human, macaque, rat, and mouse. Among datasets from all four of these tissues, genes that increased expression with age showed low conservation (IELC). Furthermore, we found that a shared trend of up-regulation among all nine tested tissues predicted lower evolutionary conservation. These results are consistent with the MA hypothesis. In addition, our functional analysis suggests that processes known to underlie senescent phenotypes in multiple tissues, apoptosis and inflammation^17^, are particularly influenced by age-dependent mutational load. Nevertheless, the methodology depends on expression data and if the function of a gene is modulated through other mechanisms, such as post-translational modifications or alterations in the interaction partners, these will not be captured in our study.

Two notes of caution are warranted. First, ADICT and IELC were not consistently detected in muscle, heart, kidney, skin, and spleen datasets, and the reason is unclear. These cases could represent false negatives due to experimental artefacts, or they might reflect heterogeneity of the ageing process. It is notable that we observed ADICT and/or IELC in only one of the 10 muscle datasets. This result could be related to a weaker signature of ageing in muscle; indeed, we could only identify age-associated genes in 4 / 10 muscle datasets (at *q* < 0.10), in stark contrast to brain datasets (18 / 19). This latter result cannot be trivially explained by reduced statistical power in the muscle data sets (sample sizes are comparable with those of brain), but could be due to increased inter-individual variation^26^ in muscle ageing. The MA hypothesis assumes late age-specific mutations; if age-specific expression patterns are absent in a tissue, we would not expect the contribution of the MA process to that tissue’s ageing. This could explain variation among tissues in ADICT prevalence.

Second, the IELC propensity could be related to up-regulation of genes evolving under positive selection, such as immune-related genes, instead of genes under weak negative selection. Hence, low conservation might reflect the accumulation of beneficial substitutions rather than weakly deleterious ones. This scenario would be consistent with the antagonistic pleiotropy (AP) hypothesis, which argues that substitutions that are positively selected for their early life benefits may be harmful in late life^27^. However, (a) our analysis is based on an estimate of negative selection rather than raw ω, and thus is not expected to be affected by positive selection; (b) when we removed genes with ω > 1 or immune genes from our analysis, we still found the same ADICT and IELC signals (Supplementary Fig. 10); (c) we compared IELC genes and 370 genes identified to be under positive selection in humans through multiple genome scans^28^, and we did not find more overlap than expected compared to the background set of all genes we analysed (Fisher’s exact test *p* = 0.3). While IELC may still partly derive from as-yet undetected positive selection in early life and represent a case of antagonistic pleiotropy, it remains at least as likely that deficient purifying selection and accumulation of slightly deleterious substitutions^29^, as predicted by the MA hypothesis, underlies the observed IELC signal.

Might the IELC phenomenon we detect contribute to physiological decline in ageing? Our finding that inflammation and apoptosis are shared functional characteristics of IELC genes in four tissues is telling, especially given increasing appreciation of the role of inflammaging, *i.e.* low-level inflammation observed in many ageing tissues^16,17,30^. There are multiple examples of how chronic inflammation can impair housekeeping functions, especially in the brain (*e.g.* refs. ^17,31^). Meanwhile, apoptosis is crucial for eliminating senescent cells during healthy ageing, and disruptions in apoptosis could lead to accumulation of dysfunctional cells over time. Conversely, apoptosis is also thought to have a role in neurodegenerative disease aetiology, such as Alzheimer’s Disease aetiology, by causing neuronal loss^32^. Our results suggest that genes involved in cellular and tissue level damage response, such as those with roles in inflammation and apoptosis, are subject to weaker purifying selection than other genes, possibly due to their limited activity early in life. The resulting mutational load leads to suboptimal regulation and function during ageing in particular tissues, when these genes show elevated activity. The MA process may thus contribute to mammalian senescent phenotypes, although at varying levels in different tissues.

## Methods

### Normalization

Affymetrix .CEL files from 23 datasets^15,24,33–50^ were downloaded from NCBI Gene Expression Omnibus (GEO)^51,52^ with accession number and EBI Array Express^53^. These raw datasets were processed using the Bioconductor “affy” package “expresso” function^54^. The selected options for the “expresso” function were: “rma” for background correction, "quantiles" for normalization, and “medianpolish” for summarization; the procedure also includes log2 transformation^55^. Whenever raw data were not provided, the pre-processed series matrix files were downloaded from NCBI GEO; the datasets were log2 transformed and quantile normalized if deemed necessary based on inspection of the pre-processed data. RNA-seq datasets were downloaded from Genotype-Tissue Expression (GTEx)^56^. These datasets were processed using log2 transformation on the gene expression levels and quantile normalization using “preprocessCore” package in R^55^. Pre-processing steps used on the analysed datasets are presented in Supplementary Table 1. Quantile normalization was performed on full datasets (without removing low expressed genes).

### Probeset-to-gene conversion

Affymetrix probe set IDs were converted to Ensembl gene IDs using the Bioconductor "biomaRt" package^57^. We used the “useMart” function to select the dataset for the species of interest, and the “getBM” function to retrieve the Ensembl gene IDs corresponding to Affymetrix probe set IDs. We then followed two steps: (a) if one probe set corresponded to more than one Ensembl gene, we removed that probe set, (b) if >1 probe set corresponded to one Ensembl gene, we chose the probe set which had the maximum expression value across all samples in that dataset. This approach used information only from the highest expressed and best-measured transcript per gene in each dataset (in other words, we discarded information from more lowly expressed and possibly noisy transcripts in that dataset).

### Age test and age-related gene sets

Genes showing age-related changes in expression levels were identified using the Spearman correlation test. We used the R “cor.test” function using “method='Spearman'” for calculating the age-expression correlation coefficient *ρ_EA_.* The *p*-values were corrected for multiple testing using the “p.adjust” function with the “Benjamini-Hochberg (BH)” method in R, yielding *q*-values as a measure of the false discovery rate. We used the nonparametric Spearman rank correlation test to overcome several problems related to conducting meta-analysis (*e.g.,* each dataset displays unique and sometimes non-normal distributions; outliers can have large effects on data analysis). We used a *q*-value cutoff of *q* < 0.10, which is a commonly used threshold (*e.g.* refs. ^15,58^). Among 58 datasets, 21 had a low number of age-related genes (*n* < 50), so to limit type II error we did not include these datasets in analyses of age-related gene sets. Gene set sizes for age-related genes and all detected genes for all 58 datasets are shown in Supplementary Table 2. Unsurprisingly, the number of age-related genes is partially affected by sample size (at Spearman correlation test *ρ* = 0.35, *p* = 0.03), but this does not influence our ability to observe ADICT (see below).

We then defined two further categories based on the expression-age Spearman correlation coefficient (*ρ_EA_*): (a) genes that showed age-related increases, with *ρ_EA_* > 0.1 and *q* < 0.1; (b) genes that showed age-related decreases, with *ρ_EA_* < -0.1 and *q* < 0.1. In addition to the *q*-value to define our cutoffs, we also used the correlation coefficient (*ρ_EA_*). Therefore genes with small effect size that can be identified in large datasets (*i.e.* with high power) but not in small datasets will not be included in our analysis. Genes with *q* > 0.10 were considered to have no change in expression level with age. Genes with *q* < 0.10 and |*ρ_EA_*| < 0.1 were discarded from further analysis.

### ADICT

The ADICT pattern was calculated as the Spearman rank correlation between age and *ρ_EC_* in each dataset, using all genes in each dataset, and using age-related genes, if detected in that dataset. The Spearman *p*-values were corrected using the BH method as described above (across all datasets included in an analysis). Note that correlation between |*ρ_A_ρ_EC_*| and sample size across datasets were negative (*ρ_A_ρ_EC_* calculated for all genes: *ρ* = -0.48, *p* < 0.001; *ρ_A_ρ_EC_* calculated for age-related genes: *ρ* = -0.64, *p* < 0.001). This is simply because finding large correlation coefficients is unlikely with large sample sizes. However, this pattern cannot explain why we observe a consistent trend for negative *ρ_A_ρ_EC_* values (i.e. ADICT) in some tissues: *e.g.* in brain 27 / 28 datasets show a negative *ρ_A_ρ_EC_* whereas only 4 / 10 muscle datasets show a negative *ρ_A_ρ_EC_*.

### Protein sequence conservation metrics

We used several types of metrics to estimate negative selection pressure on protein coding sequences.

First, we used ω_0_, a statistic based on coding sequence alignments across mammalian species. ω_0_ is estimated for the Homininae branch for human and the Murinae branch for mouse, using the branch-site model^22^. In the branch-site model, the branch of interest (the "foreground branch") is permitted to have a different distribution of *dN/dS* values than the other branches in the phylogenetic tree (the “background” branches), which are constrained to have the same distribution of *dN/dS* value among sites. The branch-site model thus estimates positive or negative selection pressure on a protein coding gene sequence. Here, we used the *dN/dS* ratio calculated for sites determined to be under negative selection. Thus, ω_0_ is expected to be a measure of the strength of negative selection on a gene. The values, calculated for each Ensembl gene, were downloaded from the Selectome database^59^.

This measure of ω_0_ can vary among genes due to multiple factors that are not the focus of this study^18^. To disentangle the effects of such factors from the effect of protein sequence conservation *per se,* we used information on GC content, CDS length, intron length, intron number, mean expression, median expression, maximum expression, tissue specificity, network connectivity, phyletic age, and number of paralogs, which were directly obtained from the Supplemental Material of Kryuchkova-Mostacci and Robinson-Rechavi (2015)^18^. To remove the effect of these variables from ω_0_, we used the “lm” function in the R “stats” package to calculate the residuals (ω_0_^*^) from a multiple regression model with ω_0_ as the response variable and all other variables as predictors. The ω_0_^*^ statistic was calculated separately for human and for mouse ω_0_ values. We used the human ω_0_^*^ data in analyses involving primate transcriptome datasets, and the mouse ω_0_^*^ data in analyses involving rodent transcriptome datasets.

Second, we calculated conservation in protein coding regions between pairs of species separated by different evolutionary distances, using *dN* (nonsynonymous substitution rate) and *dS* (synonymous substitution rate) statistics downloaded from Ensembl Biomart (v.83)^60^. Here we used “one-to-one orthologs” between human-mouse, human-elephant, and human-cow, in order to identify whether evolutionary distance between species affects estimated levels of sequence conservation. Because *dN/dS* ratios measure both the strength of negative and of positive selection, we repeated our analysis only using genes with *dN/dS* < 1 (*i.e.,* excluding the genes most likely to evolve under positive selection). In addition, GO categories and subcategories related to immune system genes (“GO:0002376”), which are known to be fast-evolving, were selected using the R “get” function. We then repeated the analysis after discarding these genes.

Finally, we calculated the conservation of protein coding sequences using the PhastCons scores (phastcons100way) downloaded from the UCSC database^25^. Phastcons100way scores each base of the human genome based on the alignment of 99 vertebrate genomes to human. To find coding regions for each gene, we used the coding start and end positions from Ensembl Biomart (v.83), combining all isoforms per gene. We obtained a list of all PhastCons scores (phastcons100way) for the coding bases of each human gene via BEDTools61 software, and then calculated the mean PhastCons score value as a metric to represent conservation of that gene’s coding region.

### Regulatory region conservation metrics

To calculate conservation for 3'-UTRs of mammalian genes, we first retrieved start and end positions of human gene 3'-UTRs from Ensembl Biomart (v.83). Due to alternative splicing, one gene may be transcribed into multiple isoforms, leading to more than one 3’-UTR per gene (which may overlap). Thus, for each gene we selected all bases annotated as part of any isoform’s 3’UTR. To calculate conservation levels of human gene promoter regions, we defined promoters as the 2000 bp upstream and downstream of a gene transcription start site (TSS), which we again obtained from Ensembl Biomart (v.83). For genes with multiple transcription start sites, we selected all bases that were located in promoter regions. To overcome possible biases that may arise from the inclusion of conserved exon regions into the regulatory region boundaries, we discarded exonic regions within the 2000 bp window around gene TSSs.

Using the BEDTools^61^ software package we obtained a list of all PhastCons scores (phastcons100way) for the defined 3’UTR bases or promoter bases of each gene. We then calculated the mean PhastCons score value as a metric to represent that gene’s 3’-UTR or promoter region conservation.

### Bootstrapping

Bootstrapping was performed using the “sample” function in R, with “replacement=TRUE”. We used bootstrapping to calculate 95% confidence intervals for the mean conservation metric among genes that showed (a) age-related increases in expression levels, (b) age-related decreases in expression levels, and (c) no age-related changes in expression levels. For each case we resampled genes 1000 times, and calculated the mean. To visually compare the conservation metric among datasets, we then subtracted the median for genes that showed no age-related change, from genes that showed age-related increase or age-related decrease. The upper and lower 2.5% quantiles are plotted in Fig. 3.

### Testing linear changes

To determine whether the relationship between individual age and *ρ_EC_* (calculated across age-related genes) was linear across adulthood, we compared linear regression models and quadratic regression models for each dataset, with *ρ_EC_* as the response variable and age as the explanatory variable (using the R “lm” function).

### Defining IELC and DEHC gene sets

We developed a non-parametric statistic, *z*, which captures the relative correlation coefficient between gene expression and age, and relative conservation level of a gene simultaneously:

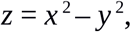

where *x* is the rank of a correlation coefficient between gene’s expression level across all detected genes in a dataset and age, and *y* is the rank of that gene’s conservation metric. Using squared values gives additional weight to differences between higher ranks. High values of *z* indicate genes that have relatively high expression and low conservation, whereas low values of *z* indicate genes that have relatively low expression and high conservation. After sorting *z* values, the top 10% of genes were included in the increasing expression and low conservation (IELC) gene set and the bottom 10% were included in the decreasing expression and high conservation (DEHC) gene set.

### Gene Ontology analysis

Here we sought to find functional groups associated with either IELC or DEHC patterns that were shared across datasets of a tissue, and across all datasets. We conducted Gene Ontology (GO) analyses for 3 GO domains - Biological Process (BP), Cellular Component (CC), and Molecular Function (MF). For this, we (I) chose GO groups showing enrichment tendencies in each dataset, using liberal cutoffs, (ii) determined the overlap among chosen GO groups among datasets, (iii) tested significance of the overlaps using random permutations of individual age in each dataset. Specifically, in each dataset, we chose GO groups with an odds ratio >1, comparing either IELC or DEHC genes (10% tails of the z statistic described above) to the rest (90%). We preferred not to use a p-value cutoff and to use liberal odds ratio cutoff (>1) in order to avoid type II error and to ensure that datasets with different numbers of genes contributed equally to downstream analysis. We then counted the number of overlapping GO groups that were thus chosen (odds ratio > 1) across all 25 datasets, or among different datasets for the same tissues. Next, we randomized ages of individuals in each dataset by conducting 1000 permutations, calculated expression correlations with age, and repeated the GO analysis using these correlation values. We finally compared the number of GO groups that showed enrichment tendency (odds ratio > 1) in the random permutations, with the observed values.

## Acknowledgements

We thank Ö. Gökçümen, M. Robinson-Rechavi, and R.Ö. Taşkent for helpful suggestions on the manuscript, all members of the METU Comparative and Evolutionary Biology Group for discussions and support. The work was supported by a Science Academy (Turkey) BAGEP Award to M.S., by the Scientific and Technological Research Council of Turkey (TÜBİTAK, projects no. 114C040 and 215Z495), and by METU BAP.

## Author contributions

M.S. and P.K. conceived the study. M.S. supervised the study. Z.G.T. and P.P. analyzed the data with contributions by H.M.D. and M.S. M.S. and Z.G.T. wrote the manuscript. All authors reviewed the manuscript.

## Competing interests

The authors declare no competing financial interests.

## Supplemental Information

### Supplementary Tables

**Supplementary Table 1**: Information about the 58 datasets used in the analysis.

**Supplementary Table 2**: Age-dependent changes in transcriptome conservation (*ρ_A_ρ_Ec_*) calculated for different conservation metrics and gene sets. **(a)** The number of age-related and all expressed genes calculated for the 58 datasets. Sheets B-F contain results (number of genes, *ρ_A_ρ_Ec_*, *p*-values) calculated using **(b)** -ω_0_ as conservation metric; **(c)** -ω_0_^*^ as conservation metric (the main result in our analyses), **(d)** -ω (or *dN/dS*) for “one-to-one orthologs” between human-mouse as conservation metric, **(e)** -ω for “one-to-one orthologs” between human-elephant as conservation metric, **(f)** -ω for “one-to-one orthologs” between human-cow as conservation metric. Sheets G-I contain results (number of genes, *ρ_A_ρ_Ec_*, *p*-values) calculated using gene sets excluding putatively positively selected genes (with ω > 1 in our data), immune system genes, and down-regulated genes in each dataset (as indicated with the “WID” suffix). We repeated this analysis using **(g)** -ω (or *dN/dS*) for “one-to-one orthologs” between human-mouse as conservation metric, **(h)** -ω for “one-to-one orthologs” between human-elephant as conservation metric, **(i)** -ω for “one-to-one orthologs” between human-cow as conservation metric. All results are calculated for all genes and age-related genes in a dataset.

**Supplementary Table 3**: Results of the comparison of the linear and quadratic regression models in 25 datasets that show an ADICT trend. A significant *p*-value indicates a better fit of the linear model, estimated by the R “lm” function.

**Supplementary Table 4**: Results of REVIGO analyses.

### Supplementary Figures

**Supplementary Figure 1.**
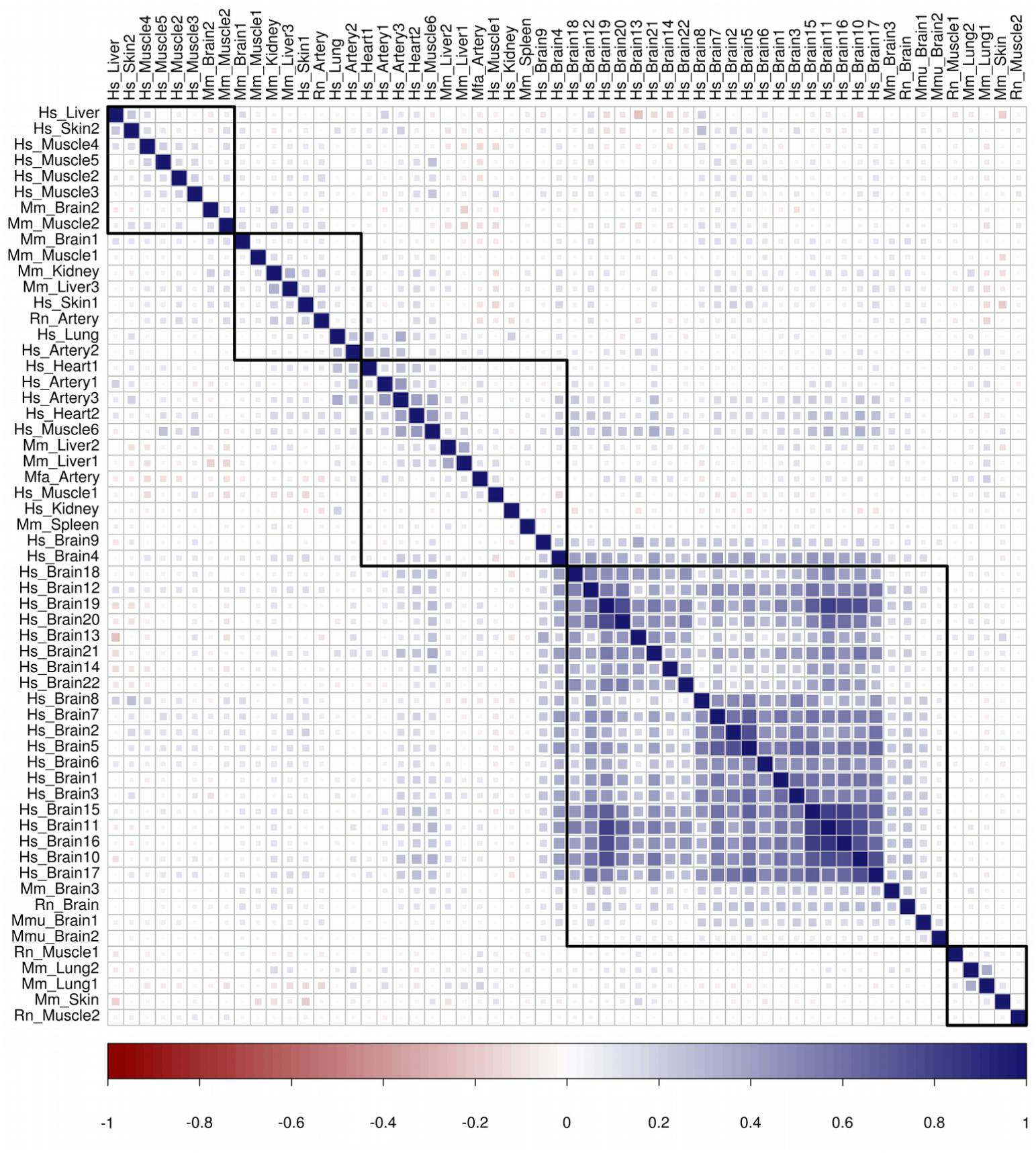
Pairwise correlations of gene-level Spearman correlation coefficients between gene expression and age (*ρ_EA_*), across all 58 datasets used in the analysis. Row and column names of the correlation matrix show each dataset, with the order determined by hierarchical clustering. Strong correlations are indicated with darker squares, red for negative and blue for positive. The number of common genes between any pair of datasets ranges from 2,516 to 21,323. The pairwise correlation coefficients across datasets range from -0.23 to 0.86. Overall there were positive correlations were found in 72% of 1653 pairwise comparisons. Among tissues belonging to the same dataset, 94% of pairwise comparisons were positively correlated among the 28 brain datasets, and 67% among the 10 muscle datasets. Among datasets belonging to the same platform, RNA sequencing and microarray datasets show 77% and 68% positive correlation among themselves, respectively. However, there is no major difference between within-platform comparisons and between-platform comparisons (Wilcoxon signed rank test *p* = 0.26).

**Supplementary Figure 2.**
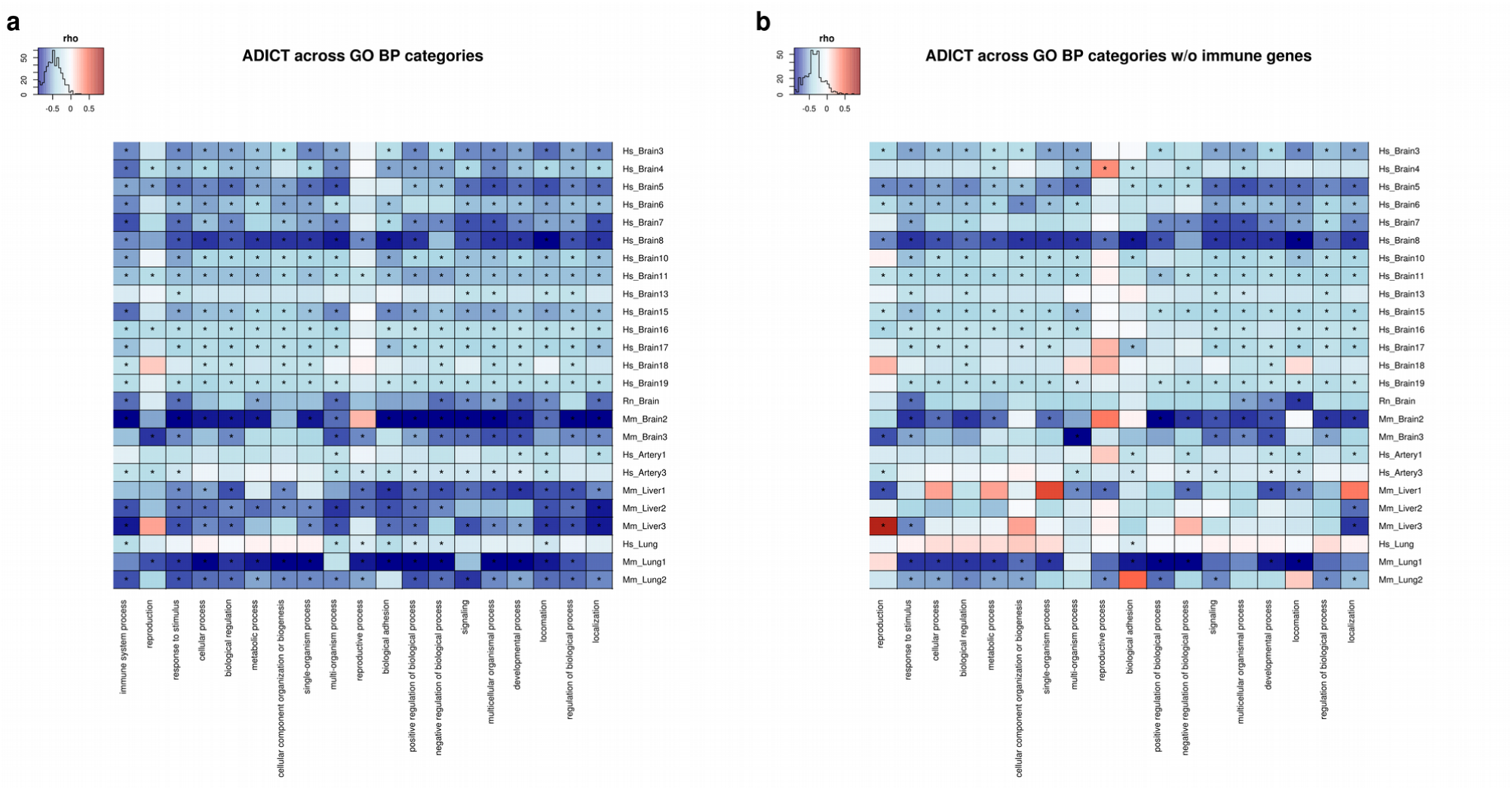
Correlation between the conservation metric (-ω_0_^*^) and expression level of genes categorized by GO BP categories that include (a) at least 1000 genes, and (b) at least 1000 genes and without immune system-related genes. These 19 GO BP categories were the only ones that had >1000 genes. Shared genes between categories were not excluded. Row and column names show each dataset and GO BP category respectively. Magnitude of Spearman correlation coefficient is indicated by the colour of the squares (darker colour shows stronger correlation): red for positive and blue for negative. The asterisks indicate, (*): *p* ≤ 0.05. Only the 25 ADICT-associated data sets are shown.

**Supplementary Figure 3.**
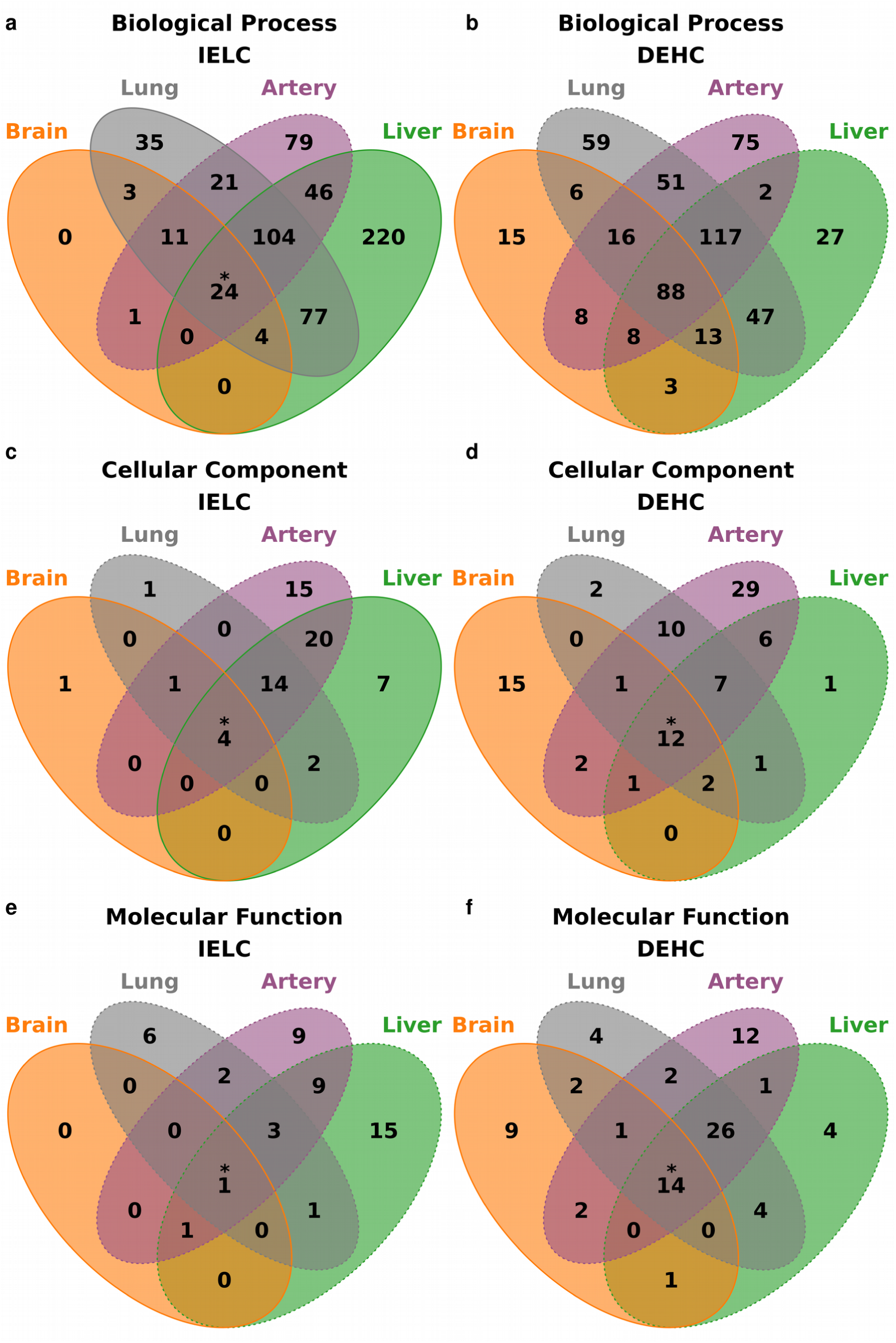
Number of GO groups enriched for genes that show increased expression with age and low conservation (IELC), and for genes that show decreased expression with age and high conservation (DEHC) across different tissues. Panels a-f show GO group enrichment results for the three GO domains. Enrichment was calculated for each tissue and for IELC or DEHC genes separately. If IELC (or DEHC) genes in a dataset showed overlap with a GO group with odds ratio > 1 (relative to other genes and other GO groups) and this was observed for all datasets of the same tissue (brain, *n* = 17; liver, *n* = 3; lung, *n* = 3; artery, *n* = 2 datasets showing ADICT), we assumed enrichment of IELC (or DEHC) genes in that GO group for that tissue (see Methods for an explanation of the rationale). The significance of the enrichment results was tested by random permutations at two levels: (a) categories found in each of the four tissues, and (b) for the categories shared between all four tissues (The asterisks indicate, (*): permutation test *p* ≤ 0.05). In panels a-f, if a tissue is demarcated by a dashed line (*e.g.* artery and lung in panel a), this indicates lack of significant GO enrichment in datasets of that tissue (*p* > 0.05). **(a)** We found 24 GO Biological Process (BP) categories (expected = 0; permutation test *p* < 0.001) enriched for IELC across all 25 datasets. Supplementary Fig. 4 contains a summary of these results as provided by REVIGO (Supek et al. 2011). Among BP GO categories, *n* = 43 (expected = 2; permutation test *p* < 0.001), *n* = 475 (expected = 93; permutation test *p* < 0.001), and *n* =279 (expected = 85; permutation test *p* = 0.002) were enriched for IELC in brain, liver, and lung, respectively. For artery, we did not find a common significant enrichment for IELC based on the permutation test. **(b)** In the 17 brain datasets, 157 GO categories (expected = 112; *p* = 0.028) were enriched for DEHC. For other tissues, we did not find a common significant enrichment for DEHC based on the permutation test. **(c)** We found four GO Cellular Component (CC) categories (“lytic vacuole”, “lysosome”, “vacuole”, “extracellular space”) enriched for IELC across all 25 datasets (expected = 0; permutation test *p* < 0.001). Among CC GO categories, *n* = 6 (expected = 0; permutation test *p* = 0.005) and *n* = 47 (expected = 20; permutation test *p* = 0.01) were enriched for IELC in brain and lung, respectively. For artery and lung, we did not find a common significant enrichment for IELC based on the permutation test. **(d)** We found *n* = 12 GO Cellular Component categories enriched for DEHC across all 25 datasets (expected = 6; permutation test *p* = 0.043). Supplementary Fig. 4 contains a summary of these results. In the brain datasets, 33 GO categories (expected = 9; *p* < 0.001) were enriched for DEHC. For other tissues, we did not find a common significant enrichment for DEHC based on the permutation test. **(e)** We found *n* = 1 GO Molecular Function (MF) category (“protein homodimerization activity”) enriched for IELC across all 25 datasets (expected = 0; permutation test *p* = 0.022). In the brain datasets, *n* = 2 GO categories (expected = 0; *p* = 0.05) were enriched for IELC. For other tissues, we did not find a common significant enrichment for IELC based on the permutation test. **(f)** We found *n* = 14 GO Molecular Function (MF) categories enriched for DEHC across all 25 datasets (expected = 5; permutation test *p* = 0.006). Supplementary Fig. 4 contains a summary of these results. In the brain datasets, *n* = 29 GO categories (expected = 8; *p* < 0.001) were enriched for DEHC. For other tissues, we did not find a common significant enrichment for DEHC based on the permutation test.

**Supplementary Figure 4.**
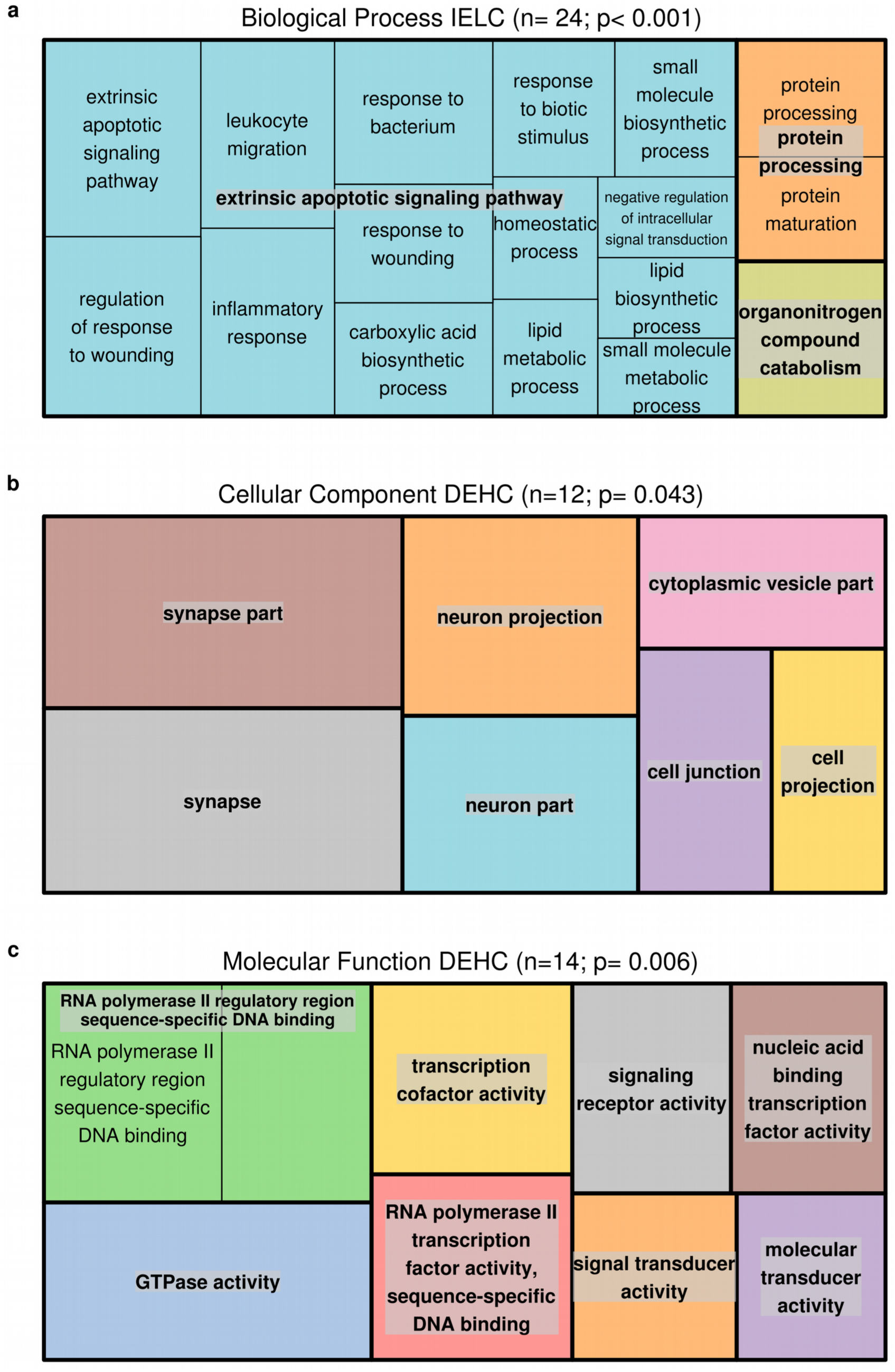
Summary of GO groups shared in all 25 datasets showing ADICT, produced using the REVIGO software (Supek et al. 2011). GO categories were selected as enriched for IELH and DEHC genes, if they showed an odds ratio > 1 in each of the 25 datasets (relative to other genes in that dataset and genes in other GO categories). Clusters are shown by rectangles and superclusters are separated by colour. **(a)** The 24 GO BP categories enriched for IELC across datasets. **(b)** The 12 GO CC categories enriched for DEHC across datasets. **(c)** The 14 GO MF categories enriched for DEHC across datasets (Supplementary Table 4).

**Supplementary Figure 5.**
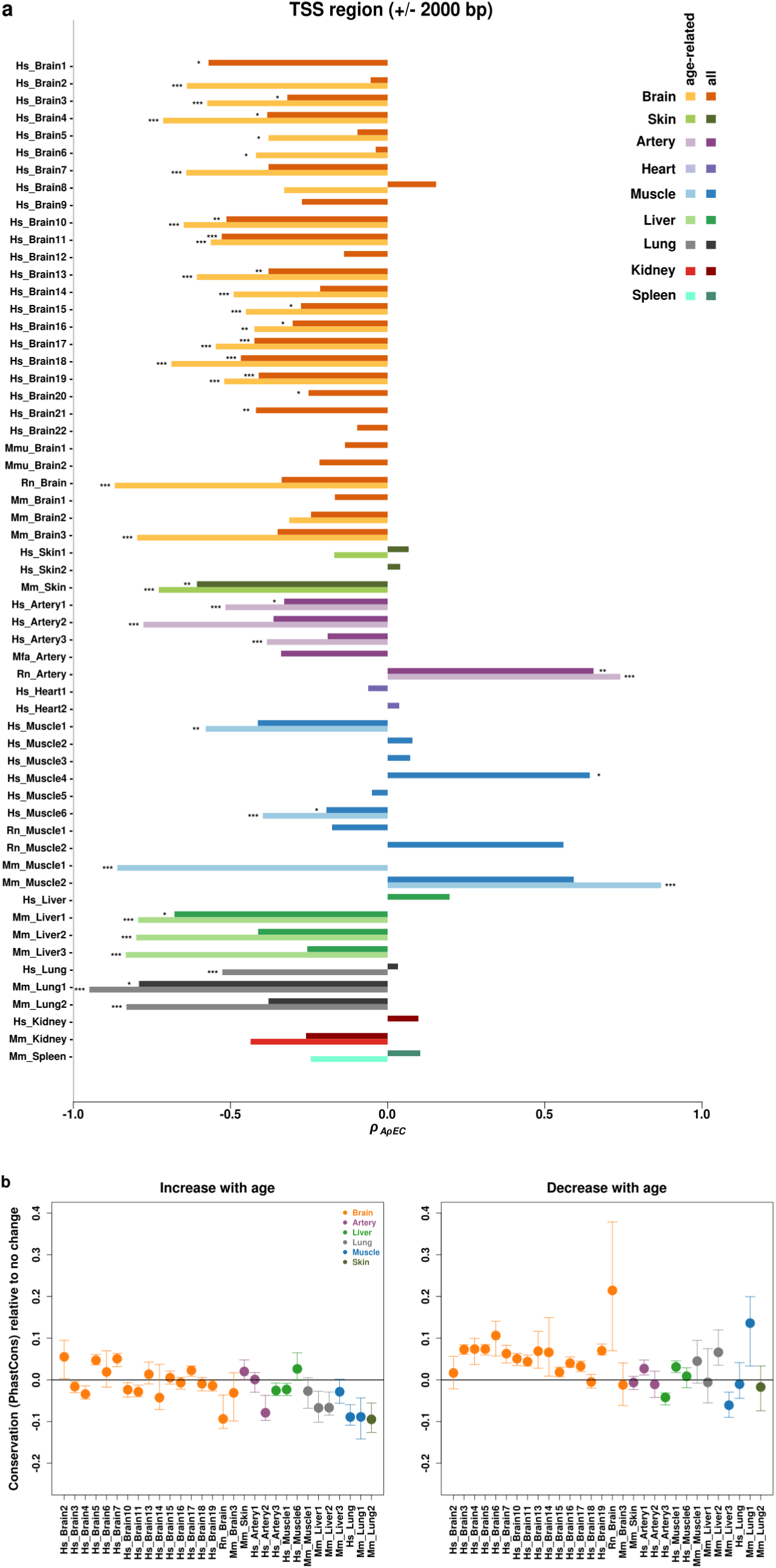
Changes in the transcription start site (TSS) region (+/- 2000 bp) conservation (PhastCons) during ageing. **(a)** The x-axis shows age-dependent change in expression level—regulatory region conservation correlation, measured by the Spearman correlation coefficient *ρ*. The results were calculated separately for each dataset, and for significant age-related genes in that dataset (light bars), as well as for all expressed genes (dark bars). Note that in 21 of 58 datasets (cases where the light bar is missing), no significant age-related gene set could be identified at *q* < 0.10. The asterisks indicate, (*): *p* ≤ 0.05, (**): *p* ≤ 0.01, (***): *p* ≤ 0.001. **(b)** Comparison of the conservation metric among gene sets showing different age-related expression level change patterns. The plots show mean conservation metric for genes showing age-related increase (left) and age-related decrease (right) in expression level, compared to mean conservation metric among genes showing no significant age-related change in expression level. The error bars indicate 95% confidence intervals calculated by 1,000 bootstraps. In 14 datasets (bootstrap support >95%), genes that show an increase in expression with age had lower regulatory region conservation, on average, than genes with no change. In 18 datasets (bootstrap support >95%) genes that show a decrease in expression with age also had higher conservation levels, on average, than genes with no change.

**Supplementary Figure 6.**
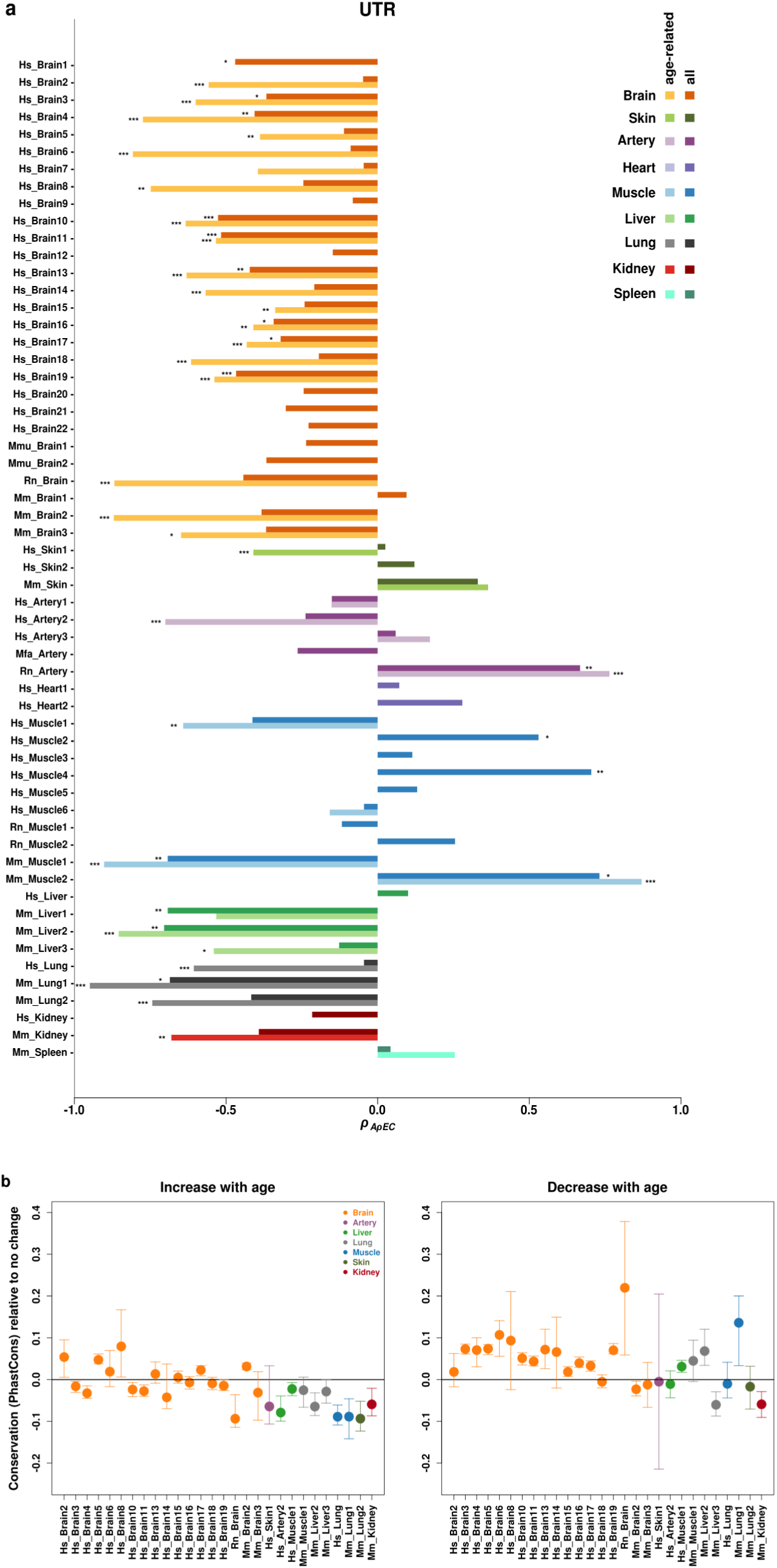
Changes in 3’ untranslated region (3’-UTR) conservation (PhastCons) during ageing. (a) The x-axis shows age-dependent change in expression level—regulatory region conservation correlation, measured by the Spearman correlation coefficient *ρ*. The results were calculated separately for each dataset, and for significant age-related genes in that dataset (light bars), as well as for all expressed genes (dark bars). Note that in 21 of 58 datasets (cases where the light bar is missing), no significant age-related gene set could be identified at *q* < 0.10. The asterisks indicate, (*): *p* ≤ 0.05, (**): *p* ≤ 0.01, (***): *p* ≤ 0.001. (b) Comparison of the conservation metric among gene sets showing different age-related expression level change patterns. The plots show mean conservation metric for genes showing age-related increase (left) and age-related decrease (right) in expression level, compared to mean conservation metric among genes showing no significant age-related change in expression level. The error bars indicate 95% confidence intervals calculated by 1,000 bootstraps. In 14 datasets (bootstrap support >95%), genes that show an increase in expression with age had lower regulatory region conservation, on average, than genes with no change. In 15 datasets (bootstrap support >95%) genes that show a decrease in expression with age also had higher conservation levels, on average, than genes with no change.

**Supplementary Figure 7.**
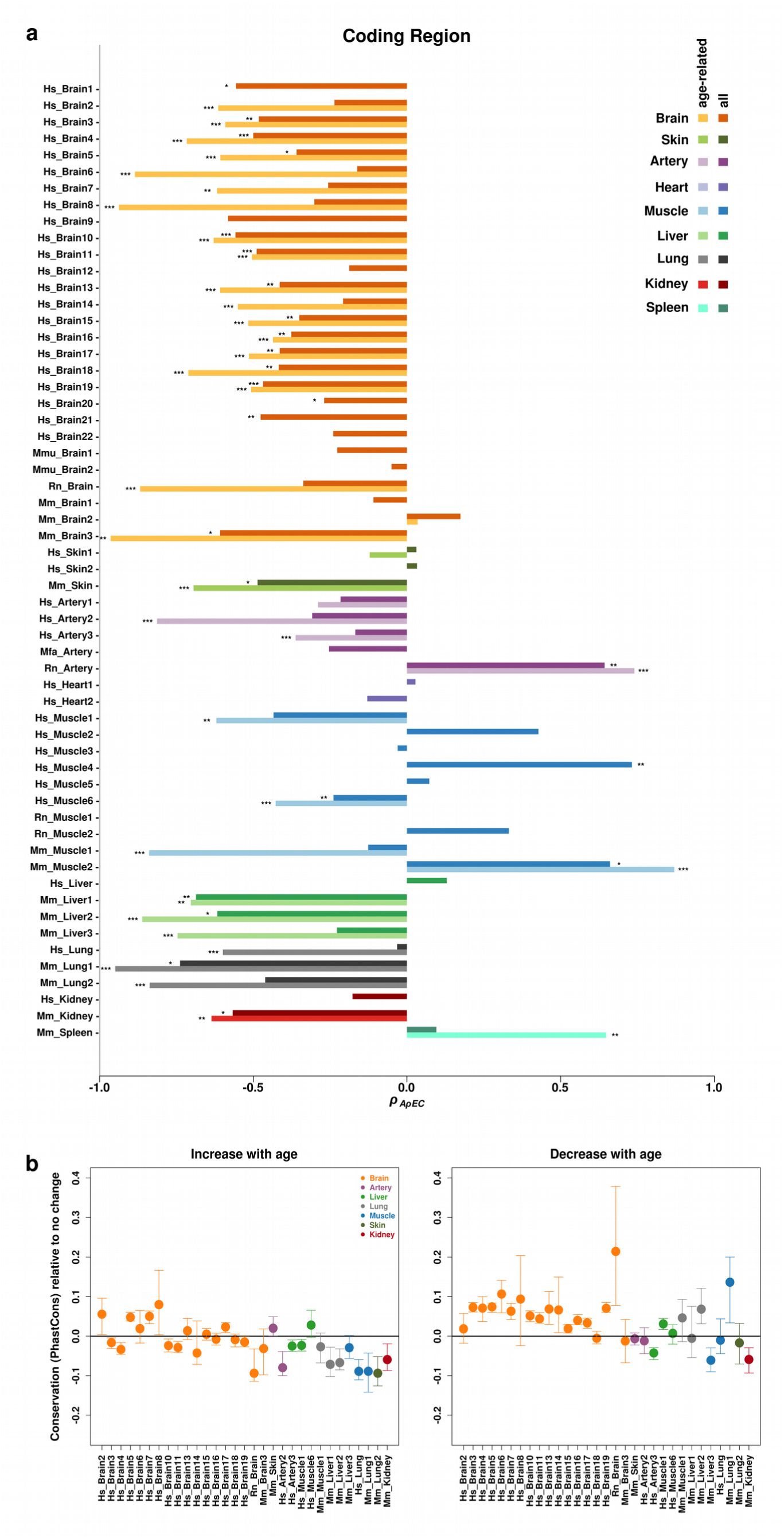
Age-related changes in conservation of coding regions measured using the PhastCons metric. **(a)** The x-axis shows age-dependent change in expression level—regulatory region conservation correlation, measured by the Spearman correlation coefficient *ρ*. The results were calculated separately for each dataset, and for significant age-related genes in that dataset (light bars), as well as for all expressed genes (dark bars). Note that in 21 of 58 datasets (cases where the light bar is missing), no significant age-related gene set could be identified at *q* < 0.10. The asterisks indicate, (*): *p* ≤ 0.05, (**): *p* ≤ 0.01, (***): *p* ≤ 0.001. **(b)** Comparison of the conservation metric among gene sets showing different age-related expression level change patterns. The plots show mean conservation metric for genes showing age-related increase (left) and age-related decrease (right) in expression level, compared to mean conservation metric among genes showing no significant age-related change in expression level. The error bars indicate 95% confidence intervals calculated by 1,000 bootstraps. In 15 datasets (bootstrap support >95%), genes that show an increase in expression with age had lower regulatory region conservation, on average, than genes with no change. In 17 datasets (bootstrap support >95%) genes that show a decrease in expression with age also had higher conservation levels, on average, than genes with no change.

**Supplementary Figure 8.**
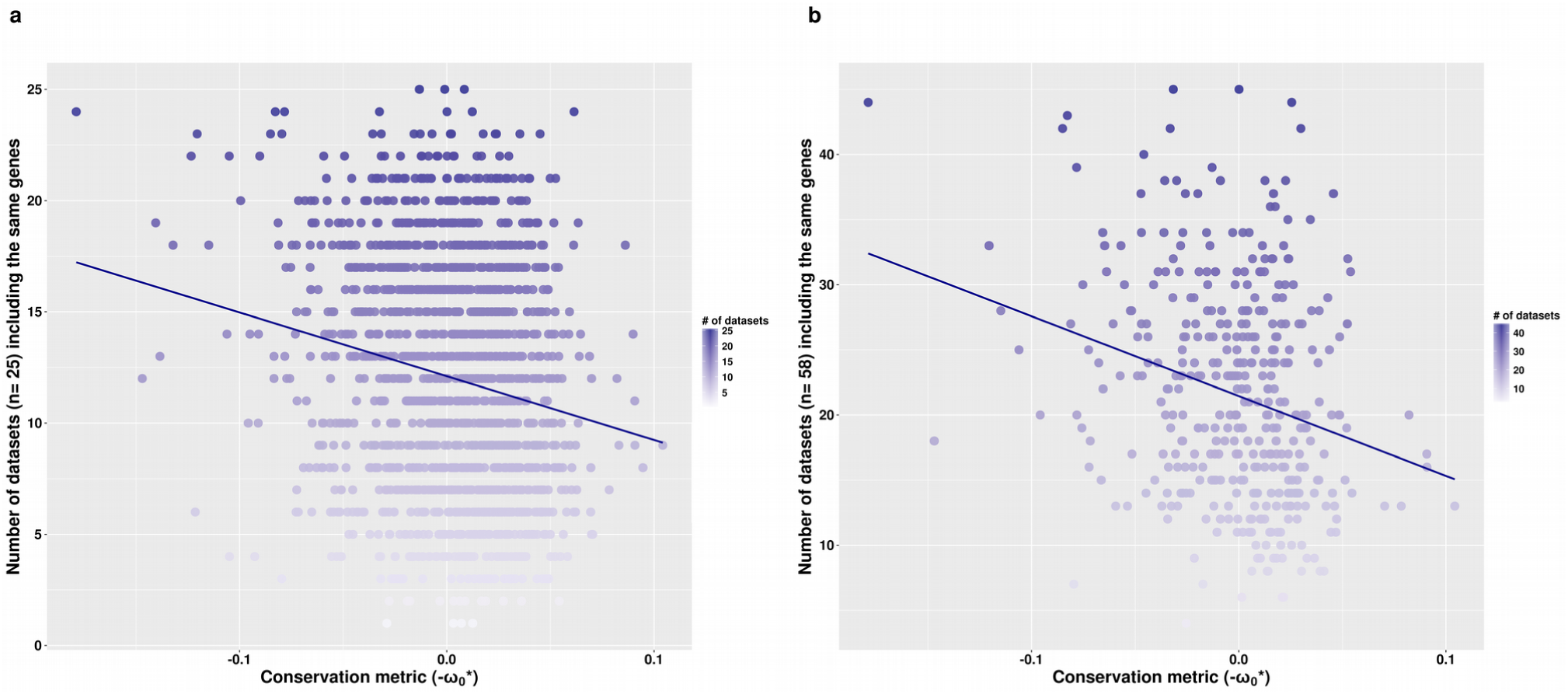
Correlation between gene protein sequence conservation metrics and the frequency of datasets where the same gene shows increase in expression with age. The x-axes and y-axes show conservation (-ω_0_^*^) and number of datasets in which a gene shows *ρ_AE_* > 0 among **(a)** the 25 ADICT-associated datasets and **(b)** all 58 datasets, respectively. Darker colour indicates higher rates of gene sharing between datasets. Spearman correlation coefficient rho is -0.17 (*p* < 0.001) for panel (a) and -0.24 (*p* < 0.001) for panel (b). Note that genes in panel (a) will not be represented in panel (b) if they are not detected in some of the 33 datasets not showing ADICT.

**Supplementary Figure 9.**
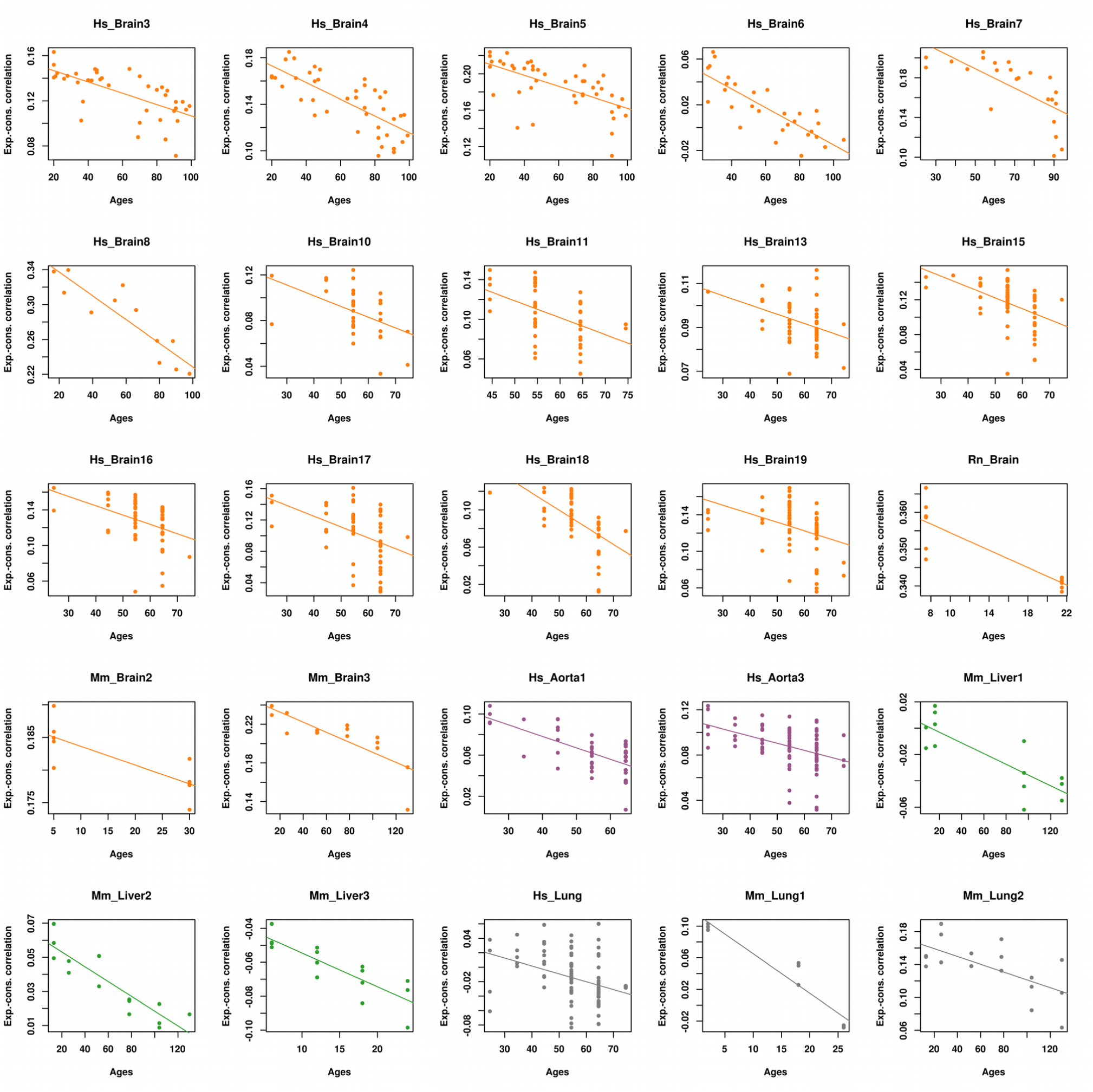
Changes in the correlation between expression and conservation metric (-ω_0_^*^) with age across all 25 datasets showing ADICT. The correlations are those represented in Fig. 2 and in Supplementary Table 2. Note that a linear model was found to fit the data better (*p* < 0.05) than an alternative quadratic model in 76% of cases, using the R function “lm”.

**Supplementary Figure 10.**
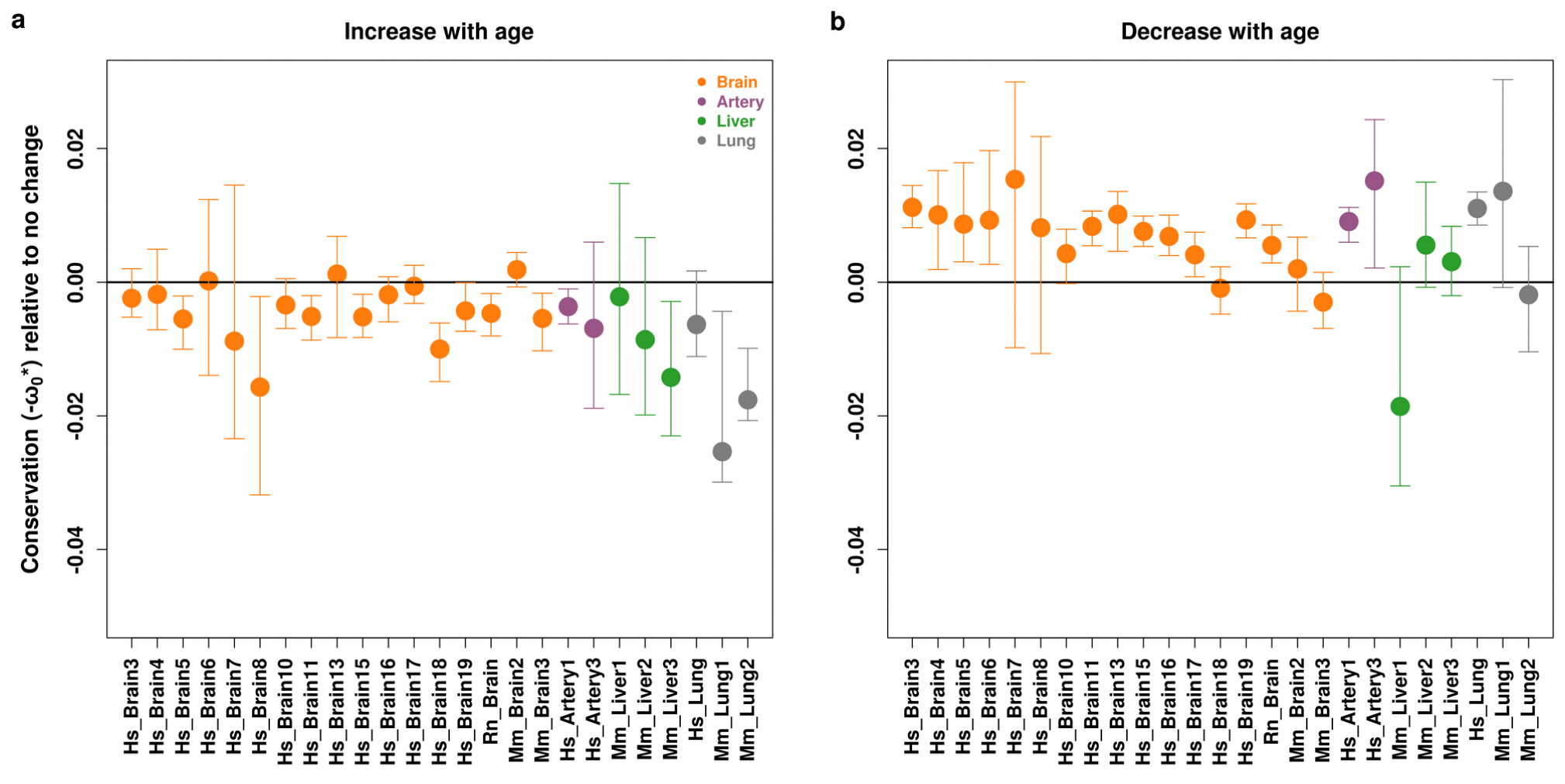
Comparison of conservation metric among gene sets that show different age-related changes in expression, after removing immune system related genes. The plots show mean conservation metric for genes showing age-related increase (left) and age-related decrease (right) in expression level, compared to mean conservation metric among genes showing no significant age-related change in expression level. The error bars indicate 95% confidence intervals calculated by 1,000 bootstraps. In 12 datasets (bootstrap support >95%), genes that show increases in expression with age had on average lower regulatory region conservation, and in 15 (bootstrap support >95%) of these datasets, genes that show decreases in expression with age also had on average higher conservation than genes with no change.

